# Local adaptation to abiotic and biotic stresses and phenotypic selection on flowering time in annual *Brachypodium spp*. along an aridity gradient

**DOI:** 10.1101/783779

**Authors:** Shira Penner, Yuval Sapir

**Affiliations:** The Botanical Garden, School of Plant Sciences and Food Security, Tel Aviv University, Ramat Aviv, Tel Aviv 69978 Israel

**Keywords:** annual grass, drought escape, life history, Mediterranean, natural selection, phenology, phenotypic cline.

## Abstract

- Plants have diverse strategies to cope with stress, including early flowering to “escape” abiotic stress and late flowering to mitigate biotic stress. Plants are usually exposed to multiple stresses simultaneously, but little is known about the impact of multiple co-occurring stresses on plant evolution.
- We tested for adaptation to both aridity and interspecific competition of the model plant *Brachypodium* spp., collected along the aridity gradient in Israel. We recorded flowering time and estimated fitness in a controlled watering experiment, with treatments mimicking Mediterranean and arid precipitation, and in two common gardens located in the extremes of the gradient (i.e., desert and mesic Mediterranean). At the latter we also manipulated interspecific competition to examine the combined effect of competition and aridity.
- Plants from arid environments always flowered earlier, but we found no selection on flowering time in the watering experiment. In the common gardens, however, the direction of selection on flowering time differed between sites and competition treatments.
- We conclude that interactions between aridity and competition drive local adaptation of *Brachypodium* in the Eastern Mediterranean basin. Variation in flowering time is an important adaptive mechanism to aridity and multiple selection agents can have interactive effects on the evolution of this trait.

## Introduction

Plants have developed diverse strategies to mitigate stress, such as early flowering to “escape” abiotic stress and late flowering to mitigate biotic stress (Ludlow, 1989; Aronson *et al*., 1993; Kigel *et al*., 2011). Plants growing naturally along environmental gradients provide a natural experiment for testing the hypothesis that natural selection leads to local adaptation. Climatic gradients are especially useful in replacing space with time to detect adaptation to different climates and potential mitigation of the response to climate changes (Etterson & Shaw, 2001; Rysavy *et al*., 2014; Rysavy *et al*., 2016). Nonetheless, climatic (abiotic) stresses are not affecting plants exclusively; biotic stresses, such as inter- and intraspecific competition also vary along climatic gradients and join abiotic stresses as selection agents (Seifan *et al*., 2010; Rysavy *et al*., 2016).

Plants have developed diverse responses to different stresses, based on highly complex mechanisms, such as changes at the developmental, transcriptome and physiological levels (Kreps *et al*., 2002; Ben Rejeb *et al*., 2014; Pandey *et al*., 2015). In addition, it has been claimed that plants respond differently to single or multiple simultaneous stresses (Rizhsky *et al*., 2004; Mittler, 2006; Mittler & Blumwald, 2010). While experimental studies usually test for the effect of a single stress, wild plant populations are usually exposed to a combination of biotic and abiotic stresses simultaneously (Ramegowda & Senthil-Kumar, 2015). Under combined stresses, plants exhibit complex physiological and molecular responses, which cannot be understood by directly extrapolating the results from studies where each stress is applied independently (Ramegowda & Senthil-Kumar, 2015). The simultaneous occurrence of biotic and abiotic stresses can cause either a negative (i.e., susceptibility) or positive (i.e., tolerance) effect on plants, depending on the species involved, its developmental stage and the intensity and duration of each stress (Tippmann *et al*., 2006; Ramegowda & Senthil-Kumar, 2015). Despite the need to understand the tolerance of plants to simultaneous biotic and abiotic stresses, there is a shortage of studies addressing this issue. To fill this gap, we examined the combined effect of drought (abiotic) and competition (biotic) stresses on the adaptation and selection of plants.

Plant adaptation to drought stress involves a change in both phenological and physiological traits, which can be categorized into three main strategies: 1) dehydration tolerance, in which plants are able to survive under decreased precipitation; 2) dehydration avoidance, which is the prevention of tissue dehydration by increasing water uptake or by decreasing water loss; 3) drought escape, which is achieved by modifying the phenology and completing all life cycles in the comfortable (short) growing season, when humidity is high (Ludlow, 1989; McKay, 2003; Sherrard *et al*., 2006). Plant responses to drought can occur on two time scales. Short-term responses include phenotypic plasticity, which is limited to the single generation and is limited in its ability to cope with the changing environment (Jump & Peñuelas, 2005; Anderson *et al*., 2012). Migration via seeds or pollen dispersal is probably too slow to track the changes already threatening many species due to climate change and decreased water availability (Jump & Peñuelas, 2005). In the long term, the evolutionary response based on standing genetic variation and driven by strong selection of heritable traits is the most promising process enabling plants to cope with the changing environment (Barrett & Schluter, 2008; Matuszewski *et al*., 2015). In this regard, the potential of a population to adapt to changes in climate will be at least partially governed by a species’ life history, namely, generation time and time span to reproduction, which occurs most rapidly in annual plants (Jump & Peñuelas, 2005; Anderson *et al*., 2012).

Aridity gradients provide natural experiments to test preadaptation to drought and reduced water availability (Petrů & Tielbörger, 2008; Lampei & Tielbörger, 2010; Tielbörger *et al*., 2010). Plant populations that have already experienced climatic stress at the drier and hotter end of the gradient may hint at phenotypic and phenological changes due to future increased aridity resulting from climate change (Holzapfel *et al*., 2006; Kreyling *et al*., 2008; Hoffmann *et al*., 2010; Kigel *et al*., 2011). The phenological shift to early flowering in annuals coping with aridity stress appears to be a widespread mechanism for adaptation to xeric environments, using the “escape” strategy to avoid stress and to reduce the risk of early senescence before seed production (Franks *et al*., 2007; Kigel *et al*., 2011). Thus, earlier flowering can be considered a preadaptation to abiotic stress (Franks *et al*., 2007; Franks & Hoffmann, 2011; Kigel *et al*., 2011).

Competition is a biotic stress constraining growth in plants (Burton, 1993). Biotic interactions may play a major role in determining the plant community structure in xeric ecosystems (Gross *et al*., 2013). Furthermore, theory and case studies suggest that there should be a predictable shift in the outcome of competitive interactions, such that competition prevails in less stressful conditions and facilitation dominates in more stressful ones (Bertness & Callaway, 1994; Pugnaire & Luque, 2001; Maestre *et al*., 2003; Seifan *et al*., 2010; Rysavy *et al*., 2016). Along an aridity gradient, contrasting stresses drive contrasting life history reactions towards the two ends of a rainfall gradient. While aridity stress towards the xeric end results in earlier flowering, competition stress in the rainy end results in late flowering (Kigel *et al*., 2011).

In the face of climate change, the fate of plant populations may be predicted by their adaptive evolutionary response to climatic conditions, derived from natural selection (Davis & Shaw, 2001; Parmesan & Yohe, 2003; Jump & Peñuelas, 2005). Adaptation to environmental stresses driven by natural selection varies in its intensity and direction across different environments, resulting in differences in local adaptation (Linhart & Grant, 1996; Joshi *et al*., 2001; Etterson, 2004; Kawecki & Ebert, 2004; Blanquart *et al*., 2013). Maintaining local adaptation in the face of climate change depends on the availability of genetic variation in relevant traits that will provide the raw materials for natural selection (Shaw & Etterson, 2012; Alberto *et al*., 2013; Lascoux *et al*., 2016). Here, we ask whether plants are locally adapted to the climatic conditions of their environment, and if they are – is natural selection the driving force of this adaptation? In addition, we test for the effect of simultaneous biotic and abiotic stresses on the adaptation to different climates and whether plants are capable to cope with multiple stresses, given they are already adapted to one of them. To answer these questions, we used wild populations of the annual grass *Brachypodium* spp., growing along the steep climatic gradient in Israel, from the mesic Mediterranean to the extreme desert.

Plants of the annual *Brachypodium* species complex (formerly known as one species, *B. distachyon*; Catalán *et al*., 2012) are self–fertile grasses (Vogel *et al*., 2009), distributed around the Mediterranean basin and in western Asia, with a wide ecological niche (Opanowicz *et al*., 2008; López-Alvarez *et al*., 2012), including along steep climatic gradients both in Israel (Kigel *et al*., 2011; Bareither *et al*., 2017; Penner *et al*., 2019) and in the Iberian peninsula (Manzaneda *et al*., 2012). The annual *Brachypodium* complex was recently split into three different karyotypic species (López-Alvarez *et al*., 2012; López-Alvarez *et al*., 2015): *B. distachyon* (diploid; 2n=10), *B. stacei* (diploid; 2n=20), and *B. hybridum* (allotetraploid; 2n=30) (Catalán *et al*., 2012). A study of *Brachypodium* spp. from the Iberian Peninsula showed that increased genome size by polyploidy is associated with aridity (Manzaneda *et al*., 2012), but two recent studies found no such association between genome size and aridity along the Israeli aridity gradient (Bareither *et al*., 2017; Penner *et al*., 2019). This lack of an association suggests that genome size is not a mechanism that drives adaptation to climate (Bareither *et al*., 2017; Penner *et al*., 2019). Given no genomic adaptation, and being indistinguishable morphologically in Israel (Penner *et al*., 2019), we consider all annual *Brachypodium* cytotypes in Israel as one taxon. While genome duplication is not the driver of adaptation to climate in *Brachypodium*, phenological variation correlates with aridity suggest that the adoption of an escape strategy by earlier flowering in arid conditions provides *Brachypodium* the ability to grow along steep aridity gradients (Kigel *et al*., 2011; Penner *et al*., 2019). The availability of plants from a wide natural distribution along the entire aridity gradient in Israel, together with the ease of growing this annual grass, offers an opportunity to test for selection regimes that may drive adaptation to climate and biotic interactions in populations of annual *Brachypodium*.

We use an experimental approach to evaluate the relative role of biotic (competition) and abiotic (aridity) stresses as selection agents on flowering time. We hypothesized that in the predictable Mediterranean climate with a higher plant density, competition is the main stress (Grime, 1979; Keddy & Shipley, 1989; Volis *et al*., 2002); thus, delayed flowering will allow the accumulation of biomass and height before the reproductive stage, increasing fitness due to better competing ability (Kigel *et al*., 2011; Colautti & Barrett, 2013; Jensen *et al*., 2013). Hence, competition will exert positive directional selection on flowering time in the Mediterranean climate. In contrast, in the arid region along the gradient, where water availability is the main constraining factor, we hypothesized that a faster life cycle (expressed as earlier flowering) will be advantageous because it will contribute to drought escape (Franks *et al*., 2007; Franks & Hoffmann, 2011; Kigel *et al*., 2011; Kooyers, 2015). Therefore, aridity, or reduced watering conditions, will exert negative directional selection on flowering time, regardless of competition. To test these hypotheses, we used plants of annual *Brachypodium* from populations along the Israeli aridity gradient and employed a common garden approach, combined with a controlled watering experiment, to measure phenotypic selection on flowering time.

## Materials and Methods

### Plant material

Seeds of annual *Brachypodium* spp. plants were collected from 20 populations in May 2013. Sites were chosen to represent the full range of the aridity gradient in Israel, from the Hermon Mountain in the North (mean annual precipitation >1200 mm) to Zenifim Wadi in the South (mean annual precipitation ∼50 mm; Table 1; **Fig. 1**). Annual *Brachypodium* plants are highly abundant throughout the Mediterranean climate region in Israel (Y. Sapir, unpublished). Collection sites were selected to evenly represent the Mediterranean region by sampling at least one population in each 100-mm rain interval (**Fig. 1a**).

**Figure 1.**
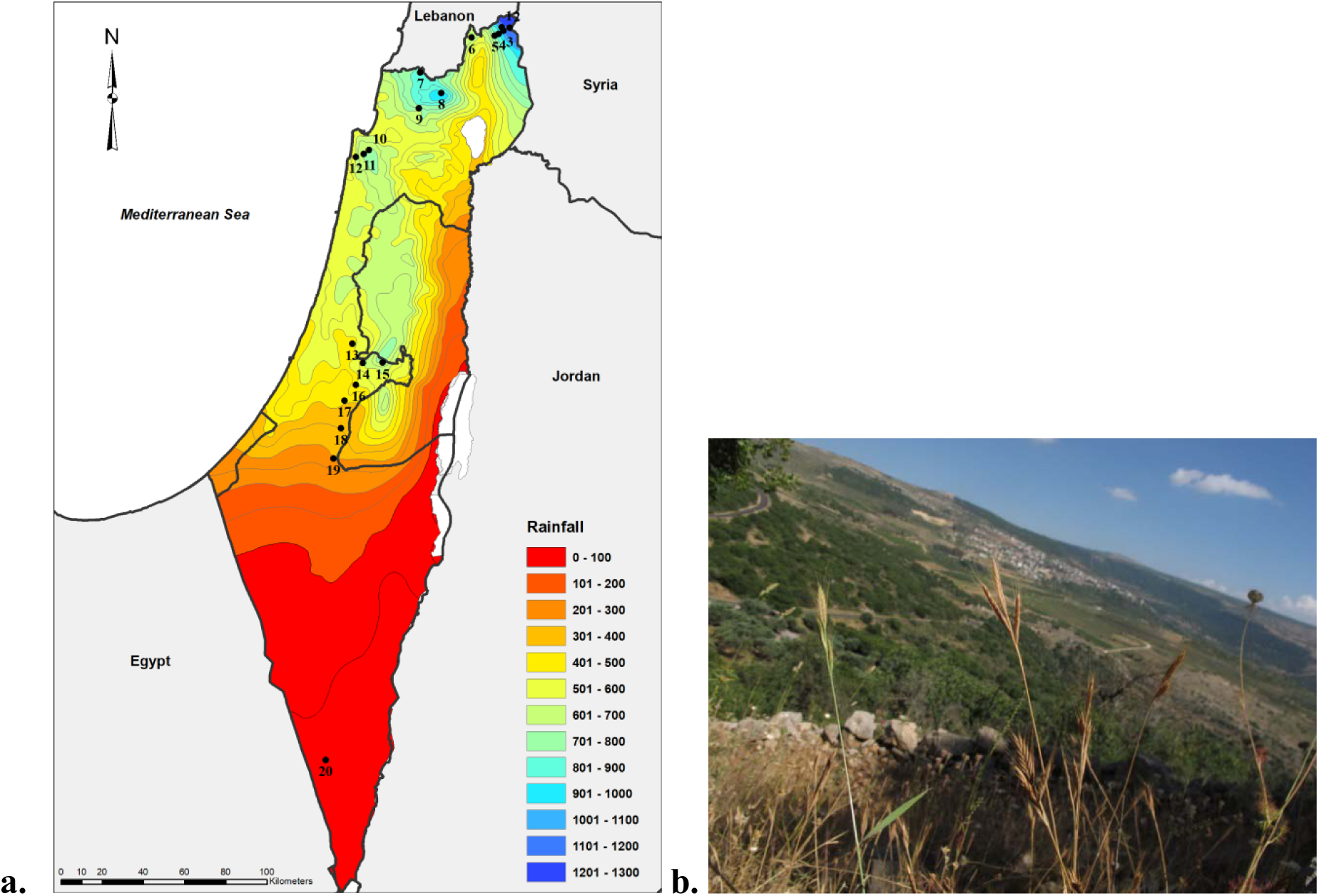
A. Geographical origin of annual *Brachypodium spp*. accessions selected in the study, on the background of the annual mean precipitation. See Table 1 for detailed information for each site. Precipitation data is the mean for years 1981-2010 (Israel Meteorological Service; http://ims.gov.il). B. *Brachypodium spp*. plants in their natural habitat in Nimrod Fortress (897 mm).

**Table 1.** Populations of *Brachypodium spp*. collected for this study. Bd21 – plants of the model *Brachypodium distachyon* accession. **Alt**. – altitude in m above sea level; **Rain** – mean annual precipitation; **Temp** – annual mean temperature; **AI** – aridity Index (mean annual precipitation/mean annual temperature); **n (Water exp.)** – number of individuals in the controlled water experiment in TAUBG, “-“ denote that this population was not represented in this experiment; **n (Com. Gar.)** – number of individuals in the common garden experiment.

|  | Site | Coordinates | Alt. | Rain (mm) | Temp (°C) | AI | n (Water exp.) | n (Com. Gar.) |
| --- | --- | --- | --- | --- | --- | --- | --- | --- |
| 1 | Hermon – Mount Kahal | 33°16'53.36N; 35°44'2.72E | 1331 | 1,098 | 13.36 | 82.16 | - | 36 |
| 2 | Hermon – Majdal Shams | 33°16'41.59N; 35°46'26.71E | 1296 | 1,214 | 13.62 | 89.15 | 32 | 64 |
| 3 | Hermon – Neve Ativ | 33°15'56.32N; 35°44'31.35E | 969 | 1,061 | 14.99 | 70.77 | 32 | 64 |
| 4 | Hermon – Nimrod Fortress | 33°15'10.89N; 35°43'4.42E | 760 | 897 | 16.21 | 55.33 | - | 72 |
| 5 | Banias | 33°14'40.74N; 35°41'49.12E | 373 | 793 | 18.16 | 43.66 | - | 64 |
| 6 | Tel Hai | 33°14'11.97N; 35°34'31.35E | 303 | 718 | 18.02 | 39.85 | - | 64 |
| 7 | Shtula | 33° 05'4.70N; 35°18'26.59E | 631 | 805 | 16.70 | 48.21 | 32 | 64 |
| 8 | Mount Meron | 32°59'39.98N; 35°24'57.25E | 1132 | 909 | 14.25 | 63.79 | - | 52 |
| 9 | Karmiel | 32°55'40.06N; 35°18'2.56E | 280 | 782 | 17.55 | 44.56 | - | 60 |
| 10 | Carmel – Rakafot | 32°44'38.02N; 35° 2'33.75E | 510 | 701 | 16.67 | 42.05 | 32 | 64 |
| 11 | Carmel – Kedumim Quarry | 32°43'40.29N; 35° 0'52.85E | 297 | 709 | 17.77 | 39.90 | - | 64 |
| 12 | Carmel – Oren Gorge | 32°42'52.05N; 34°58'23.23E | 54 | 612 | 18.81 | 32.54 | 32 | 64 |
| 13 | Modi'in near Train Station | 31°53'45.27N; 34°57'30.09E | 140 | 493 | 19.54 | 25.23 | - | 64 |
| 14 | Kiryat Anavim | 31°48'46.63N; 35° 6'47.53E | 699 | 696 | 16.82 | 41.39 | - | 64 |
| 15 | Eshtaol Forest | 31°48'43.27N; 35° 0'45.39E | 290 | 587 | 19.01 | 30.88 | - | 56 |
| 16 | Bet-Shemesh Heights | 31°42'58.81N; 34°58'36.90E | 318 | 489 | 18.91 | 25.81 | 32 | 64 |
| 17 | Avishur Plato | 31°38'49.23N; 34°55'06.23E | 367 | 419 | 18.52 | 22.62 | 40 | 80 |
| 18 | Amatzya | 31°31'30.30N; 34°53'58.49E | 367 | 370 | 18.70 | 19.79 | 32 | 64 |
| 19 | Lahav | 31°23'31.72N; 34°51'42.90E | 421 | 294 | 18.62 | 15.79 | 32 | 64 |
| 20 | Zenifim Wadi | 30° 4'21.36N; 34°49'41.88E | 470 | 51 | 20.18 | 2.53 | 22 | 44 |
| 21 | Bd21 |  |  |  |  |  | 2 | 4 |

For climatic characterization of each population, mean annual precipitation and mean annual temperature data were retrieved from the Israeli meteorological service website (http://ims.gov.il). From these data we calculated an aridity index as the ratio between the mean annual precipitation and temperature (Manzaneda *et al*., 2012; Penner *et al*., 2019).

To reduce the effects of microscale spatial heterogeneity, seeds were collected from the same habitat in all Mediterranean sites, namely, rocky south-facing slopes. In semi-arid and desert habitats (<300 mm), populations are scattered, and thus seeds were collected where found, regardless of the exact microhabitat. In 2013, we collected one spike per plant from at least 20 haphazardly chosen plants at each site. Between 11 and 20 (mean = 16) maternal families (i.e., different plants from the same population) were grown in the Tel Aviv University Botanical Garden (TAUBG) in the fall of 2013. Seeds were germinated in a commercial germination soil mix and watered twice a day via an upper mist for 20 minutes in the TAUBG nursery. At the end of the growing season (between mid-April and early July 2014), seeds were collected from all plants and stored in paper bags at 4°C. In November 30^th^ 2014, 6 seeds from each maternal family were planted and grown under the same conditions as in 2013. Growing plants for two generations under equal and controlled conditions allows us to discard maternal effects.

### Controlled Watering experiment

To specifically test the effect of drought stress, we tested the phenology and fitness of plants from along the gradient in two contrasting watering regimes. We used four experimental plots in TAUBG. The plots were covered from the top by 2 × 4 m sheds of transparent plastic panels, in a way that exposed the plants to all climatic conditions except rain. Light intensity, measured using a light meter (EXTECH, USA), was measured under each shed at three different points (top center, middle center and bottom center of the shed). No significant difference in light intensity was found between the four sheds (P=0.07). Plants were watered from 15 December 2016 to 28 May 2017 by a dripper irrigation system, using drippers of 2 l·h^-1^ placed 30 cm apart. Each shed was randomly assigned to either of two treatments: two sheds mimicked Mediterranean conditions with an equivalent of 600 mm annual rain, achieved by watering 150 minutes per week, and the other two sheds mimicked semi-arid conditions with an equivalent of 200 mm annual rain, achieved by watering 50 minutes per week.

We chose a subsample of ten populations along the gradient for the controlled drought experiment (Table 1). Between 11 and 20 (mean = 16) maternal families of plants from each population were chosen, and 4 of the seeds from each maternal family were randomly picked for the experiment, in addition to two *Brachypodium distachyon* plants of the model accession genotype Bd21. Two seeds of each family and one of Bd21 were planted in each treatment (Table 1). Seeds were sowed directly in the ground at 20-cm intervals, in a randomized block design for each treatment. From each maternal family we sowed two seeds in each point. If both seeds germinated, we randomly removed one of the seedlings one week after germination; in this way, each maternal family was represented once in each treatment. A total of 320 plants, representing 160 maternal families, were included in this experiment.

### Common garden experiments

In November 2016 we set up two experimental common garden plots in two different climates: one in a mesic Mediterranean climate, near Tel Hai Academic College (33°14’N, 35°34’E, altitude of 280 m above sea level), with mean annual rain of 770 mm, and the second in a semi-arid climate, within the Bergman campus of Ben-Gurion University in Be’er Sheva (31°15’N, 34°47’E, altitude of 280 m above sea level), with mean annual rain of 195 mm. At each site, we set up two plots of 5 × 5 m, of which one plot had all the natural vegetation removed to prevent competition, while local annual plants were allowed to grow in the other.

At each site we planted 618 plants from 20 populations along the gradient, in addition to two *Brachypodium distachyon* plants of the model accession genotype Bd21 (Table 1). Each maternal family was represented in all plots and sites by one plant. Plants were grown at 20-cm intervals, in a fully randomized block design in both sites.

To initiate germination we watered the experimental plots immediately after sowing, but no additional watering was applied except natural rain. As in the controlled watering experiment above, two seeds were sowed in each point, and one of the two seedlings was randomly thinned one week after germination. A total of 2,472 seeds were sowed in both experimental sites, of which 1, 236 seedlings were left to grow after germination (618 in each site).

### Phenotypic measurements

Phenotypic traits measured were identical for both watering regime and common garden experiments. Traits were chosen to represent indicators of drought adaptation (Kigel *et al*., 2011; Catalán *et al*., 2012).

**Flowering time** was recorded daily for the controlled watering experiment in TAUBG, throughout the growing season, between February 27^th^ and May 28^th^ 2017. In the two common garden sites, flowering time was recorded twice a week, between March 4^th^ and May 28^th^ 2017. We defined flowering time as the number of days from sowing to first appearance of the awns of the first spike (inflorescence; see Penner *et al*., 2019).

**Morphology** was measured for all plants in May 2017, at the time of harvesting. Morphological measurements included eleven traits (determination of traits following Catalán *et al.*, 2012): (1) inflorescence length, measured on the central stem of the plant (the primary stem), from the bottom of the lowest spike to the top of the upper one (not including the awns); (2) spike length, measured as the length of the upper spike on the primary stem (excluding awns); (1) stem length, measured for the primary stem from the ground to the bottom of the lowest spike; (4) number of stems, counted as the number of tillers; (5) aboveground biomass, measured as the dry weight of the shoot, excluding the roots and spikes (because plants were harvested dry after senescence, there was no need to dry them); (6) number of nodes on the primary stem; (7) awn length, measured as the length of the longest awn of the top spike of the primary stem; (8) lemma length, measured from the base of the lowest seed on the top spike of the primary stem to the base of the awn of this flower; (9) spike length to the 4th lemma, measured as the length from the base of the lowest seed on the top spike of the primary stem to the base of the awn of the 4^th^ seed from the bottom of this spike; (10) caryopsis length, measured as the length of the lowest seed of the top spike of the primary stem; (11) upper glume length, measured in the top spike on the primary stem.

**Fitness** was assessed for both survival and reproduction. Survival was defined as the success of a plant to survive until flowering before senescence. Plants that died before flowering or did not set flowers until the end of the season were recorded as not-survived. The survival rate per population was estimated as the fraction of plants that set flowers out of all plants of that population in the same treatment. Reproductive output was evaluated as the total number of spikes of a plant. In a preliminary study, we found an average number of 6.5 seeds per spike, with relatively low variance of this number (Y. Sapir, unpublished). Thus, we used the total number of spikes as a surrogate for the seed output. Annual *Brachypodium* spp. are mainly self– fertile (Vogel *et al*., 2009), thus maternal fitness (total number of spikes) is a good estimation of life-time fitness in this plant.

### Statistical analyses

Statistical analyses were performed using R (R Development Core Team 2014). The measured morphological traits are highly correlated in *Brachypodium* (Penner *et al.*, 2019). Hence, to account for these correlations, as well as to avoid bias due to multiple tests, we collapsed all 11 morphological traits measured to a low number of dimensions using principal component analysis (PCA). For the plants in the controlled watering experiment in TAUBG, the first principal component (PC1) explained 41.3% of the variance, mainly due to traits of the reproductive organs, such as inflorescence length, spike length and 4^th^ lemma length. The second principal component (PC2) explained 12.1% of the variance, affected mainly by lemma length, upper glume length and caryopsis length. For the plants in the common gardens, PC1 explained 39.4% of the morphological variance, mainly due to plant size and reproductive traits, such as inflorescence length, stem length, spike length and number of stems. PC2 accounted for 19.8% of the morphological variance, affected mainly by reproductive-related traits, associated with seed size, such as lemma length, caryopsis length, and upper glume length. We used the values of each plant on the first principal component (PC1) in each experiment as a surrogate for morphology and size.

#### Effect of stress on phenotypic traits

Generalized linear model (GLM) was used for analyses of covariance (ANCOVA) to evaluate the changes in flowering time, morphology or fitness of populations along the gradient as a function of the climate (aridity index) in their site of origin and treatments. For the controlled watering experiment we used the following statistical model:

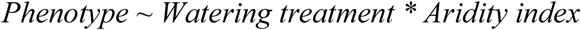

For the common gardens experiment we used the following model, with competition treatment nested within the experimental site:

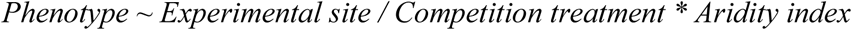

To assess the effect of shed number on phenotype in the controlled watering experiment we added shed number as a co-factor in the statistical model. Nonetheless, in the preliminary analyses this co-factor had no significant effect (P>0.05) in all models and thus was removed from the analysis.

To analyze survival (measured as the proportion of plants), we used a binomial GLM model; to analyze the number of spikes (maternal fitness), which are count data, we used the Poisson distribution. To evaluate the inclusion of interactions in the statistical models, we compared models with and without the interaction terms using the Akaike Information Criterion (AIC).

#### Selection analyses

We estimated phenotypic selection on flowering time following Lande & Arnold (1983), by calculating selection gradients (*β*). Fitness was defined as the total spike number and was relativized (mean fitness = 1) by dividing by the average number of spikes in each combination of climatic origin, site and treatment. Flowering time was standardized to the mean = 0 and variance = 1, using function “scale” in R. The use of standardized trait values and relative fitness enabled the estimation of standardized partial linear regression coefficients corresponding to the strength and direction of directional selection acting on flowering time (Lande & Arnold, 1983). Only plants that survived and set flowers were included in this selection analysis.

Because sample size in each combination of population, experimental site and treatment was small, we pooled plants from populations of similar climate in the origin site. In the controlled watering experiment, we defined the range of 50 to 400 mm as arid climate, and areas with >400 mm as Mediterranean climate. In the common garden experiment, we defined the range of 50 to 400 mm as arid climate, the range of 400 to 800 mm as Mediterranean climate, and areas with >800 mm as mesic Mediterranean climate.

We used GLM for analyses of covariance (ANCOVA) to test whether climatic origin, site and treatment affected the extent and direction of selection. To achieve this goal, we tested the significance of the covariance between flowering time and each of the three factors by including their interactions in the model. We used the following statistical model for the controlled watering experiment:

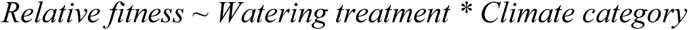

We used the following statistical model for the common gardens:

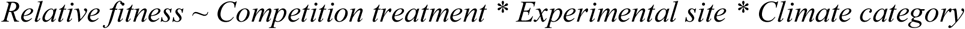

Because relative fitness is not taken from the normal distribution, we used the natural log transformation to test for significance.

## Results

### Effect of stress on phenotypic traits

#### Controlled watering experiment

##### Survival

Despite the differences between treatments in water availability, all plants survived under both treatments (Table 2). Thus, there was no point in analyzing the effect of treatment and aridity on survival.

**Table 2.** Effects of site, treatment and climatic origin on phenotypic traits in the controlled watering experiment and the common garden experiments. P (survival) – proportion of plants survived and reached flowering stage; Flowering time – days lapsed from sowing; Fitness – total number of spikes. Values are means ± standard errors for each combination of site, climatic group and treatment.

| Site | Treatment | Climate in origin | P (survival) | Flowering time | Fitness |
| --- | --- | --- | --- | --- | --- |
| Tel Aviv | Wet treatment | Arid | 1 | 88.4 ± 1.28 | 16.2 ± 1.29 |
| | | Med | 1 | 96.4 ± 1.11 | 10.6 ± 0.9 |
| | Dry treatment | Arid | 1 | 88.8 ± 1.13 | 9.2 ± 1.14 |
| | | Med | 1 | 96.4 ± 1.11 | 7.2 ± 0.9 |
| Mediterranean (Tel Hai) | Competition | Arid | 0.94 | 134.6 ± 0.78 | 2.7 ± 1.23 |
| | | Med | 0.99 | 136.5 ± 0.4 | 2.5 ± 0.59 |
| | | Wet Med | 0.98 | 139.1 ± 0.59 | 3.1 ± 0.89 |
| | No competition | Arid | 1 | 138.0 ± 0.8 | 14.8 ± 1.24 |
| | | Med | 1 | 138.6 ± 0.4 | 15.1 ± 0.59 |
| | | Wet Med | 1 | 146.6 ± 0.59 | 16.9 ± 0.9 |
| Arid (Be'er Sheva) | Competition | Arid | 0.67 | 129.6 ± 1 | 1.4 ± 3.37 |
| | | Med | 0.33 | 129.7 ± 0.46 | 1.5 ± 1.85 |
| | | Wet Med | 0.11 | 132.2 ± 0.56 | 1.9 ± 2.49 |
| | No competition | Arid | 1 | 123.5 ± 0.99 | 29.4 ± 3.31 |
| | | Med | 0.99 | 126.1 ± 0.46 | 32.1 ± 1.84 |
| | | Wet Med | 0.99 | 132.0 ± 0.55 | 35.6 ± 2.47 |

##### Flowering time

In both treatments, we found that plants from the arid climate set flowers about eight days earlier than plants from the Mediterranean or wet Mediterranean climates (Table 2; detailed results for all populations are presented in the Supporting Information Table S1). Model selection using AIC revealed that the most informative model included the main factors of the aridity index in the origin of the plant and watering treatment, without their interaction (*Flowering time ∼ Treatment* + *Aridity index*). While watering treatment had no significant effect on flowering time, aridity did have a significant positive effect on flowering time (Table 3; **Fig. 2a**).

**Figure 2.**
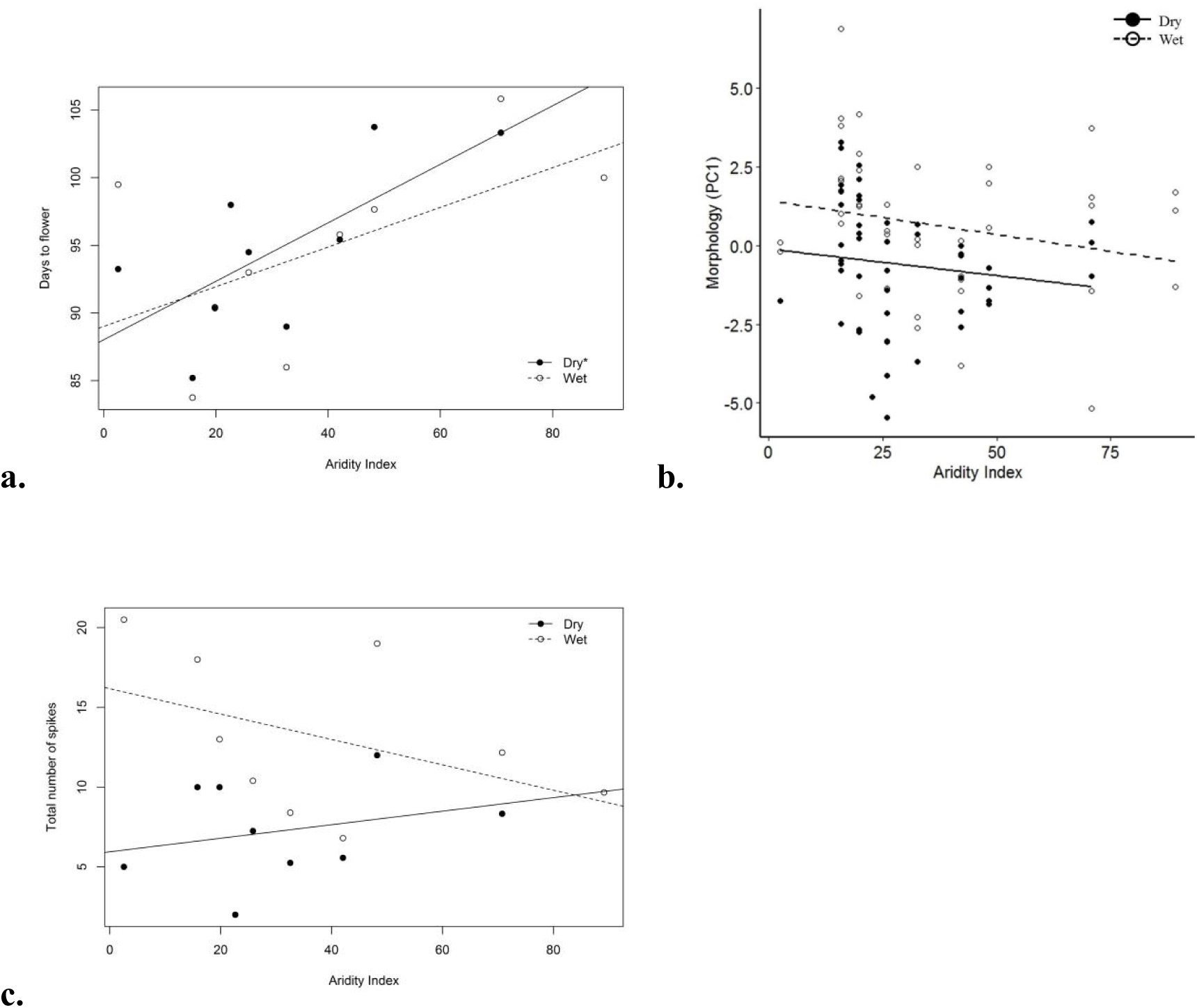
Effects of aridity and watering treatment on phenotypic traits of annual *Brachypodium spp*. in a controlled watering experiment in TAUBG. In figures A, C, points are means for populations, and in figure B, points are all the individuals who participated in the experiment. Empty circles and dashed lines denote plants in the wet treatment and full circles and solid lines denote plants in the dry treatment. Slopes significantly different from zero are denoted by an asterisk (*). A. The effect of watering treatment on flowering time; treatments not significantly affecting flowering time but affecting aridity (P=0.971, P=0.008, respectively; Table 3). B. The effect of watering treatment on morphology (PC1); treatments significantly affecting morphology and not aridity (P=0.006, P=0.085, respectively; Table 3). C. The effect of watering treatment on fitness (total number of spikes); the interaction of treatment and aridity does not significantly affect fitness (P interaction = 0.187; Table 3).

**Table 3.**
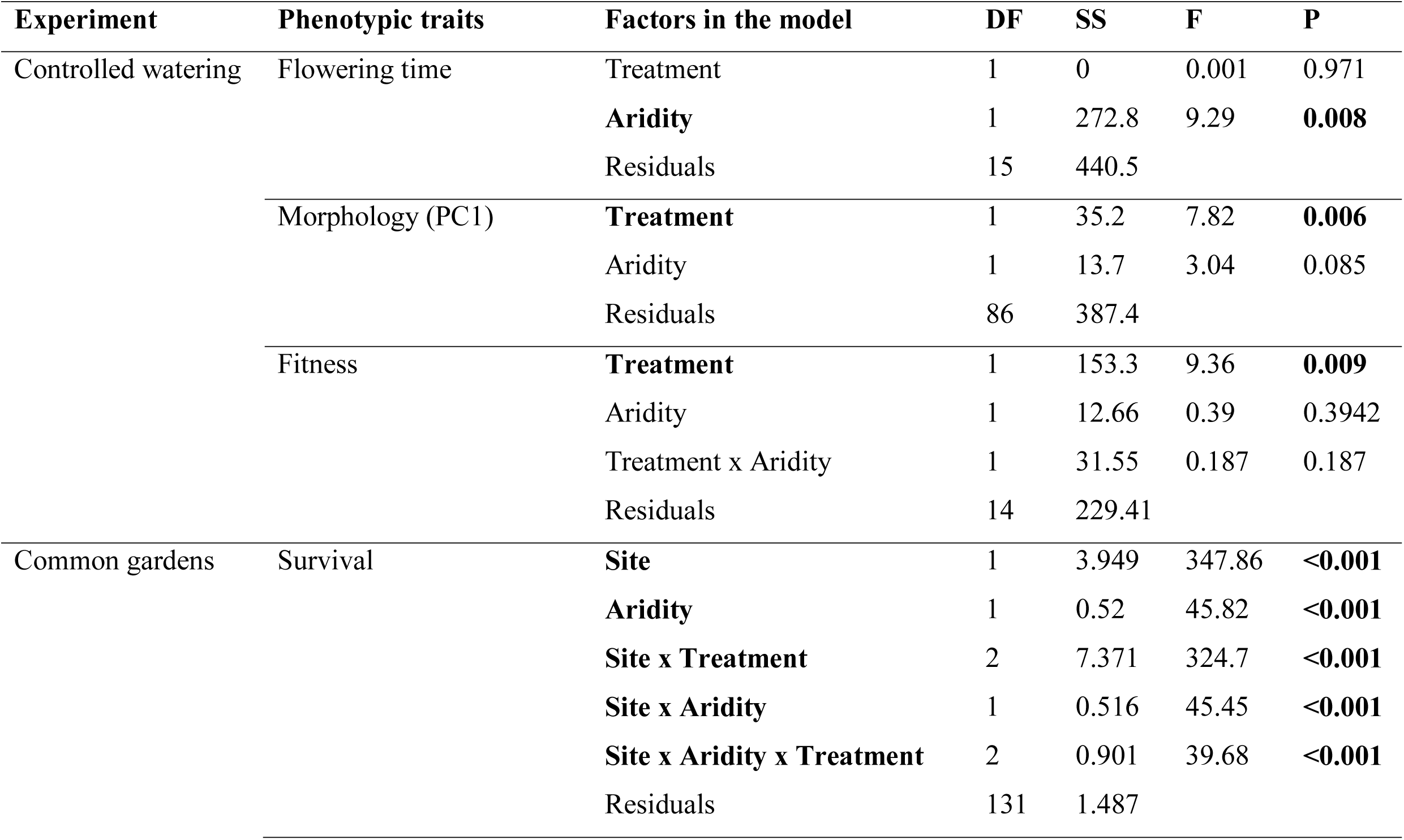
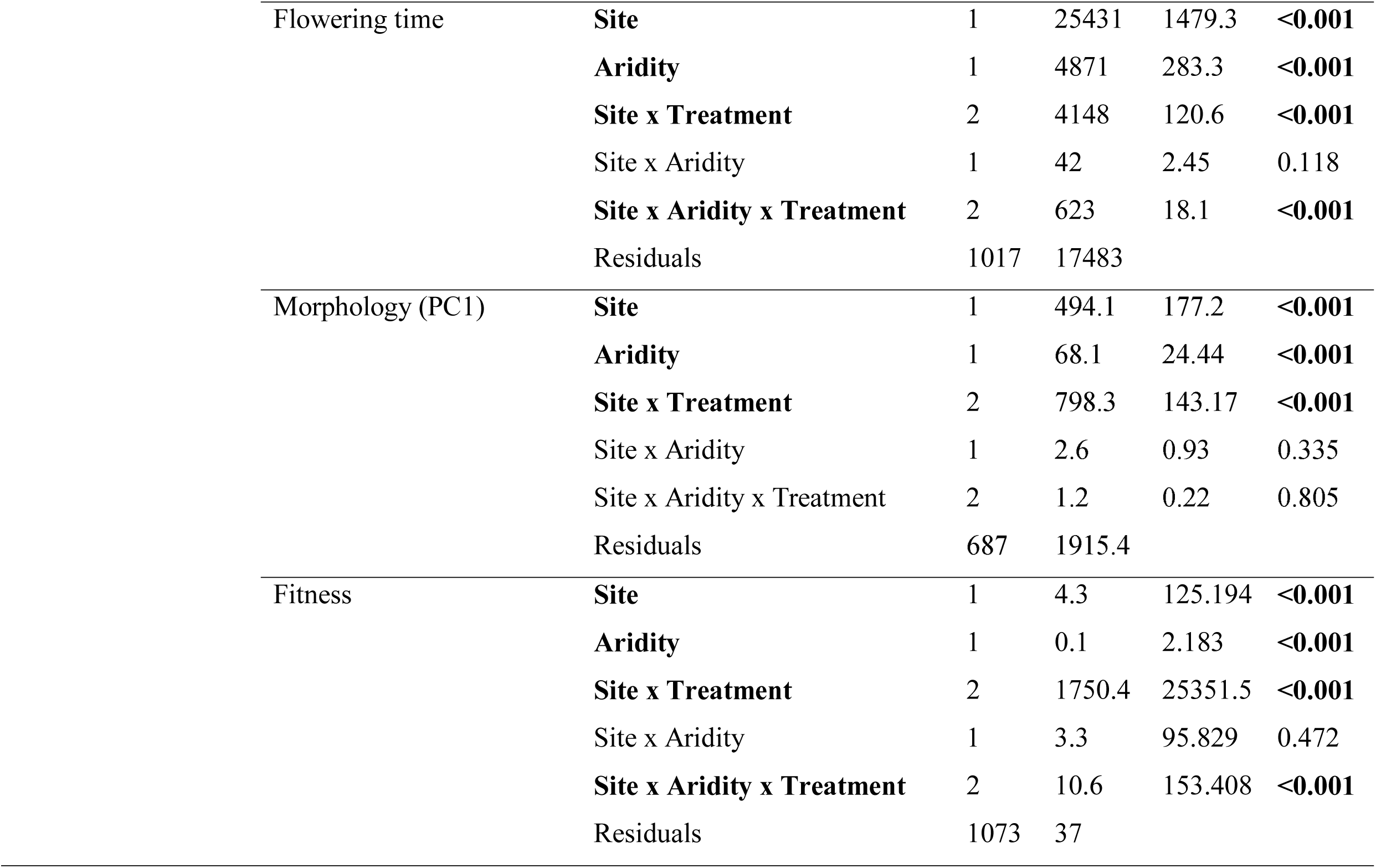
Statistical analyses of variance from both the controlled watering experiment and the common garden experiment. In the common garden experiment, treatment (competition) was nested within site. Significant effects are shown in bold. For binomial data, a binomial distribution was considered, and for count data, a Poisson distribution was considered.

##### Morphology

Plants in the wet treatment were slightly larger than in the dry treatment without an apparent effect of aridity (**Fig. 2b**). The most informative model included only the main effects of aridity and watering treatment, without their interaction (*Morphology ∼Treatment + Aridity index*). The watering treatment had a significant effect on morphology, while aridity did not (Table 3; **Fig. 2b**).

##### Fitness

The total number of spikes was higher for plants in the wet treatment compared with the dry treatment, and in each treatment, the plants from the arid climate had a greater number of spikes on average (Table 2; **Fig. 2c**). The most informative model included both main effects and their interaction (*Total number of spikes ∼ Treatment * Aridity index*). The watering treatment significantly affected fitness, while aridity index and the interaction term did not (Table 3; **Fig. 2c**).

#### Common garden experiments

##### Survival

Almost all plants in the Mediterranean site (Tel Hai) survived; in contrast, survival in the arid site (Be’er Sheva) depended on the treatment: while in the absence of competition most plants survived, survival was low with competition (Table 2; detailed results for all populations are presented in the Supporting Information Table S2; **Fig. 3a**). The most informative model included all the main effects and their interactions, with treatment nested within site (*Survival ∼ Site /Treatment * Aridity index*). The experimental site and aridity index, as well as all treatment (nested within site) and all interactions, significantly affected survival (Table 3; **Fig. 3a**).

**Figure 3.**
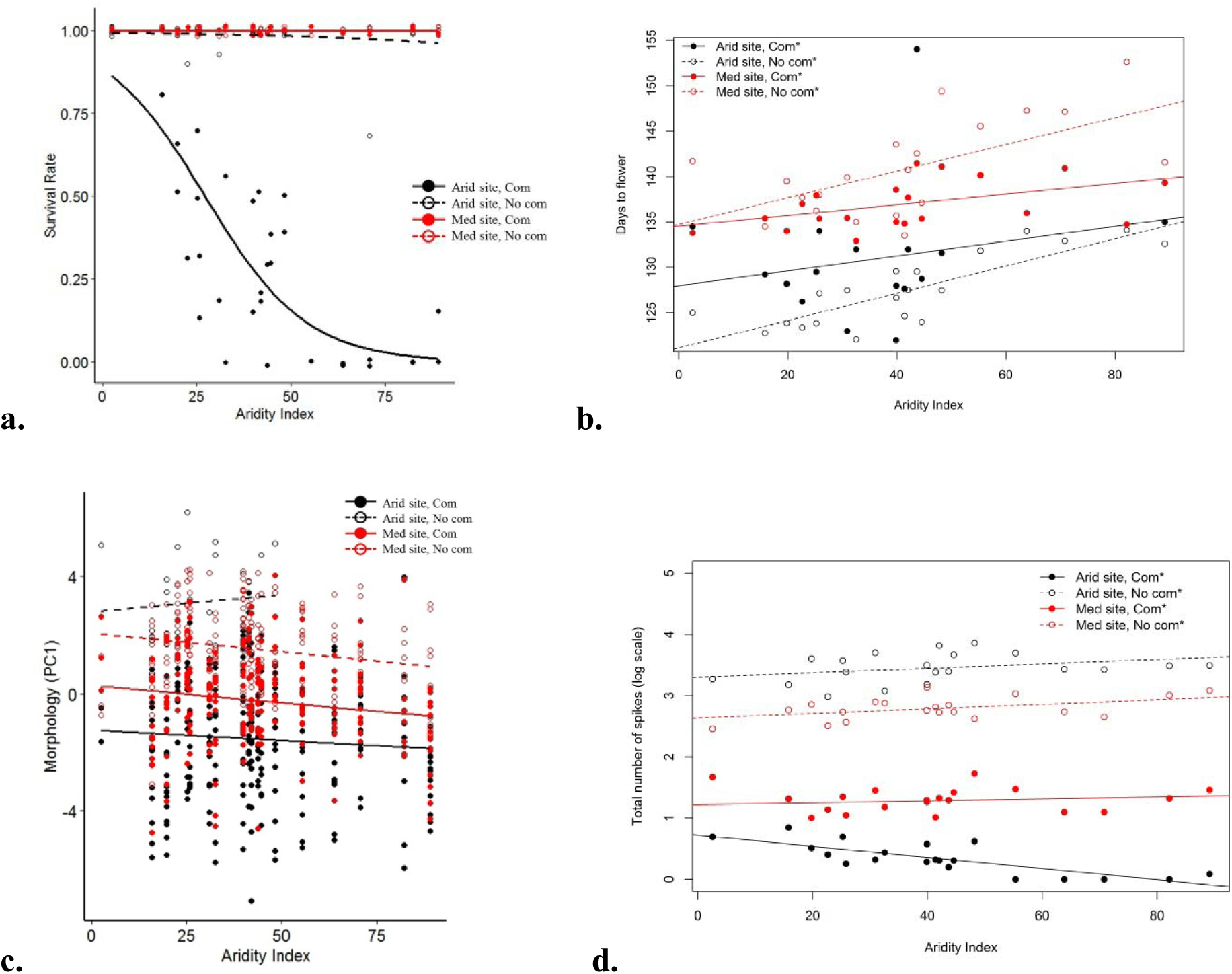
–Effect of site and treatment and their interaction in terms of phenotypic traits of *Brachypodium spp*. from populations along the aridity gradient in Israel in two common gardens. A. The effect of site*treatment on survival, the binomial smooth function was used to create a nonlinear trend line; the interaction of site, treatment and aridity significantly affect survival (P interaction < 0.001; Table 3). B. The effect of site*treatment on phenology (flowering time); the interaction of site, treatment and aridity significantly affects flowering time (P interaction < 0.001; Table 3). C. The effect of site*treatment on morphology (PC1); the interaction of site, treatment and aridity does not significantly affect morphology (P interaction = 0.805; Table 3). D. The effect of site*treatment on fitness (total number of spikelets) after ln-transformation; the interaction of site, treatment and aridity significantly affects fitness (P interaction < 0.001; Table 3). In figures A, B, D points are the means for the populations, and in figure C, points are all the individuals who participated in the experiment – empty points denote plants with no competition treatment, and full points denote plants with competition treatment. The colors represent the different sites (the arid site in black and the Mediterranean site in red). Dashed and solid lines also denote plants with and without competition treatment, respectively. Slopes significantly different from zero are denoted by asterisk (*).

##### Flowering time

Similar to the controlled watering experiment, flowering time was earlier with increased aridity, but the difference in flowering time was also affected by competition and climate: while in the arid site, flowering occurred overall earlier than in the Mediterranean site (Table 2), the gradual change in flowering time from the arid to the Mediterranean climate in the origin of the plants was steeper without competition (**Fig. 3b**). The most informative model included all the main factors and their interactions, with treatment nested within site (*flowering time ∼ site /treatment * Aridity index*). The experimental site and aridity index had a significant effect on the flowering time, while in the partial interaction terms, only the interaction between site and treatment significantly affected flowering time, and the interaction between site and aridity index did not (Table 3; **Fig. 3b**).

##### Morphology

Plants in both experimental sites were larger (i.e., higher values on the first principal component) without competition, compared to plants growing under competition treatment (**Fig. 3c**). The most informative model included all the main effects, with treatment nested within site, and the interaction (*Morphology ∼ Site /Treatment * Aridity index*). The experimental site and aridity index had a significant effect on morphology, as well as the effect of treatment nested within site, while the interaction between site and aridity index was not significant (Table 3; **Fig. 3c**).

##### Fitness

The total number of spikes was an order of magnitude larger without competition in both sites (Table 2), but did not show a strong association with aridity (**Fig. 3d**; note the logarithmic scale of the y-axis). The most informative model included all the main effects, with treatment nested within site, and the interaction (*Total number of spikes ∼ Site /Treatment * Aridity index*). The experimental site and aridity index had a significant effect on fitness, as well as the effect of competition treatment nested within site, while the interaction between site and aridity index was not significant (fitness was ln-transformed for the significance tests, Table 3).

The three-way interaction term of site x treatment x aridity index was significant (Table 3; **Fig. 3d**).

### Phenotypic selection on flowering time

#### Controlled watering experiment

Analysis of covariance revealed no significant interaction of climate origin and treatment with flowering time in their effect on fitness (Table 4). Individual selection gradients for each combination of climate origin and treatment were not significantly different from zero, except for plants from the arid climate that experienced reduced water (dry treatment), which had a significant negative directional selection (s = –0.235, P=0.050; relative fitness was ln-transformed for the significance tests; Table 5; **Fig. 4a**).

**Figure 4.**
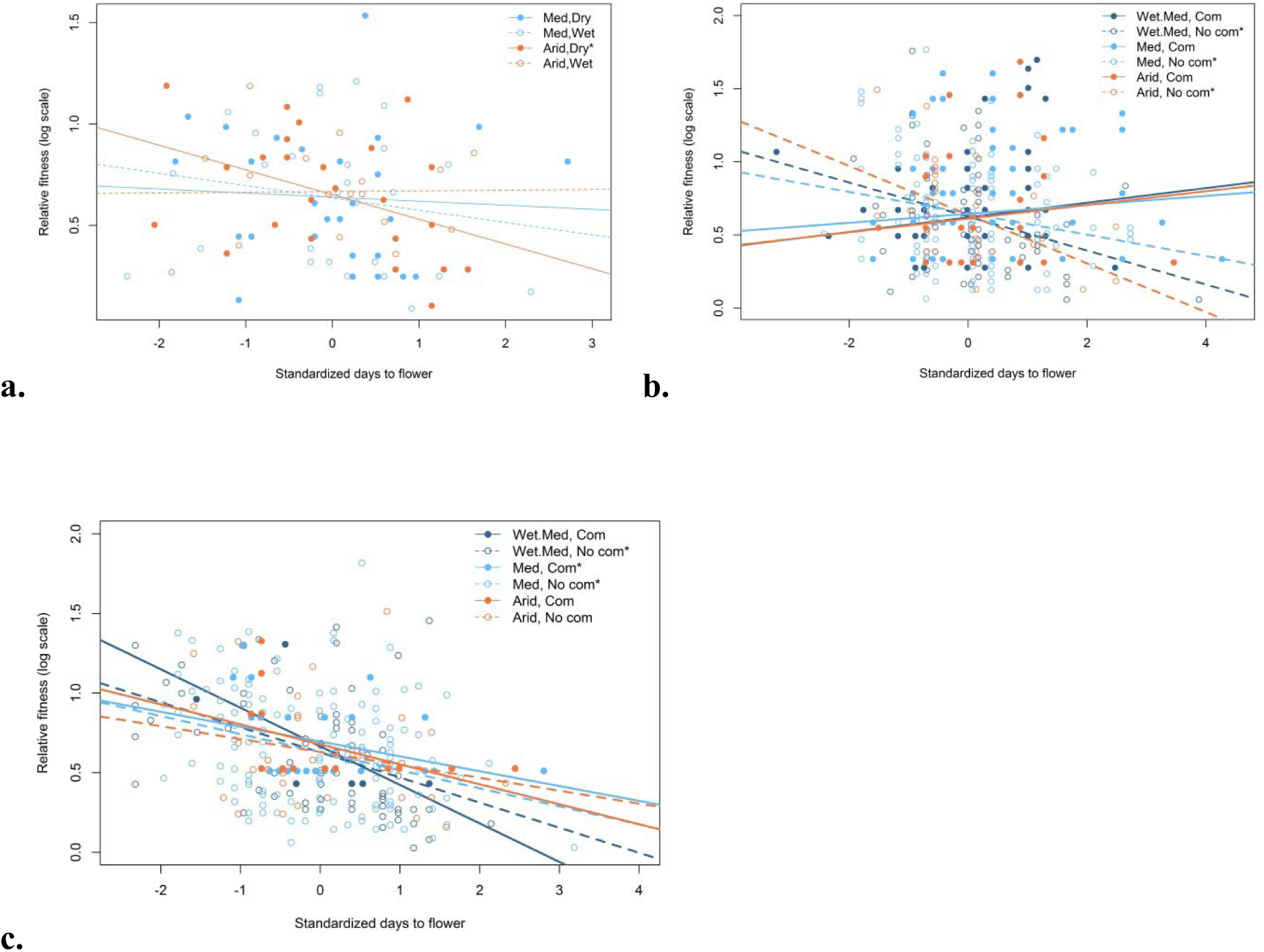
Selection on flowering time in both the controlled watering experiment and the common gardens. Selection was measured as the effect of phenology (flowering time) on relative fitness (total number of spikes) using the standardized phenotype and a logarithmic scale to normalize the relative fitness data. A. Selection on flowering time in the controlled watering experiment; the interaction of flowering time, treatment and climate does not significantly affect the relative fitness (P=0.227; Table 4). B. Selection on flowering time in the Mediterranean site in the common garden experiment; the interaction of flowering time, treatment and climate does not significantly affect the relative fitness (P=0.58; Table 4). C. Selection on flowering time in the arid site in the common garden experiment; the interaction of flowering time, treatment and climate does not significantly affect the relative fitness (P=0.74; Table 4). The colors represent the different climatic origins (Wet Med in dark blue, Med in light blue and Arid in orange). Lines denote partial regression slopes for no competition treatment (dashed line) or competition treatment (solid line). Slopes significantly different from zero are denoted by an asterisk (*). All values are the means of climatic origin*site*treatment combinations.

**Table 4.** Analyses of variance for selection analyses of flowering time. Because of the non-normal distribution of relative fitness, ln-transformation for significance testing was employed. Significant effects are in bold.

| Experiment | Factors in the model | DF | SS | F | P |
| --- | --- | --- | --- | --- | --- |
| Controlled watering | Flowering time | 1 | 0.75 | 1.78 | 0.105 |
|  | Treatment | 1 | 0 | 0.002 | 0.93 |
|  | Climate | 1 | 0.19 | 0.444 | 0.65 |
|  | Flowering time x Treatment | 1 | 0.12 | 0.276 | 0.583 |
|  | Flowering time x Climate | 1 | 0.13 | 0.297 | 0.687 |
|  | Treatment x Climate | 1 | 0.01 | 0.034 | 0.712 |
|  | Flowering time x Treatment x Climate | 1 | 0.43 | 1.007 | 0.227 |
|  | Residuals | 88 | 37.27 |  |  |
| Mediterranean common garden | **Flowering time** | 1 | 1.42 | 2.259 | **0.034** |
|  | Treatment | 1 | 0.21 | 0.334 | 0.402 |
|  | Climate | 2 | 0.17 | 0.139 | 0.79 |
|  | **Flowering time x Treatment** | 1 | 11.48 | 18.292 | **<0.001** |
|  | Flowering time x Climate | 2 | 0.87 | 0.69 | 0.564 |
|  | Treatment x Climate | 2 | 1.44 | 1.144 | 0.247 |
|  | Flowering time x Treatment x Climate | 2 | 0.6 | 0.482 | 0.58 |
|  | Residuals | 440 | 276.15 |  |  |
| Arid common garden | **Flowering time** | 1 | 13.06 | 24.508 | **<0.001** |
|  | Treatment | 1 | 0.81 | 1.519 | 0.051 |
|  | **Climate** | 2 | 2.88 | 2.699 | **0.044** |
|  | Flowering time x Treatment | 1 | 0.26 | 0.487 | 0.255 |
|  | Flowering time x Climate | 2 | 0.96 | 0.9 | 0.225 |
|  | Treatment x Climate | 2 | 0.12 | 0.117 | 0.739 |
|  | Flowering time x Treatment x Climate | 2 | 0.31 | 0.289 | 0.74 |
|  | Residuals | 282 | 150.3 |  |  |

**Table 5.** Selection on flowering time in the controlled watering and common garden experiments. Selection gradients (*β* ± standard errors) were calculated for each combination of site, climatic origin and treatment.

| Site | Climatic origin | Treatment | $\beta$ | P |
| --- | --- | --- | --- | --- |
| Tel Aviv | Arid | Wet | $0.023 \pm 0.127$ | 0.86 |
| | | Dry | $-0.235 \pm 0.114$ | **0.050** |
| | Med | Wet | $-0.102 \pm 0.138$ | 0.47 |
| | | Dry | $-0.029 \pm 0.146$ | 0.85 |
| Mediterranean (Tel Hai) | Arid | Competition | $0.158 \pm 0.172$ | 0.365 |
| | | No competition | $-0.355 \pm 0.119$ | **0.006** |
| | Med | Competition | $0.092 \pm 0.066$ | 0.166 |
| | | No competition | $-0.222 \pm 0.081$ | **0.007** |
| | Wet Med | Competition | $0.213 \pm 0.135$ | 0.122 |
| | | No competition | $-0.244 \pm 0.096$ | **0.014** |
| Arid (Be'er Sheva) | Arid | Competition | $-0.285 \pm 0.147$ | 0.073 |
| | | No competition | $-0.161 \pm 0.147$ | 0.283 |
| | Med | Competition | $-0.21 \pm 0.096$ | **0.037** |
| | | No competition | $-0.226 \pm 0.065$ | **0.001** |
| | Wet Med | Competition | $-0.524 \pm 0.367$ | 0.226 |
| | | No competition | $-0.297 \pm 0.085$ | **0.001** |

#### Common garden experiments

Analyses of covariance revealed a significant effect of flowering time on fitness (that is, flowering time is under selection overall; Table 4). In the Mediterranean site (Tel Hai), competition treatment interacted significantly with flowering time, suggesting different selection regimes under competition and without competition (Table 4). Indeed, while all plants under competition experienced positive directional selection on flowering time, without competition this selection was negative (**Fig. 4b**). Only the negative directional selection gradients for plants without competition were significantly different from zero (Table 5).

In the arid site (Be’er Sheva), the climate in the origin site from which plants were collected (either arid, Mediterranean, or wet Mediterranean) also significantly affected fitness (Table 4). Nonetheless, none of the interactions were significant (Table 4), providing no evidence for differential selection of climate in origin or competition on flowering time. In all combinations of climate origin and competition treatment, negative selection on flowering time occurred in the arid site (**Fig. 4c**), but the selection gradient was significant only for plants from the Mediterranean climate in both treatments and for plants from the wet Mediterranean climate without competition (Table 5).

## Discussion

In this research, we used an experimental approach to evaluate the relative role of biotic (competition) and abiotic (aridity) stresses and their synergistic effect on fitness and on a life history trait (flowering time), in addition to their roles as selection agents that drive local adaptation. We found that competition interacted with aridity and local adaptation in terms of their effect on survival, where plants from the Mediterranean climate survived less under competition in the arid experimental site (**Fig. 3a**). Furthermore, flowering time occurred earlier for plants from arid climates compared with plants from Mediterranean climates, in both arid and Mediterranean climates or under high or low watering regimes (**Fig. 2a, 3b**). Interestingly, competition influenced the rate of change in flowering time with aridity in the origin of plants, and this gradual change was steeper without competition (**Fig. 3b**). As expected, plants were larger and had higher reproductive output without stress (biotic or abiotic; **Fig. 2b** and **2c**, **Fig. 3c** and **3d**). Finally, while no selection on flowering time was observed in the controlled watering experiment (**Fig. 4a**), in both common gardens (Mediterranean and arid) we found differential selection on flowering time, where in the Mediterranean site competition changed the extent and direction of selection (**Fig. 4b**). This finding suggests that aridity (abiotic) stress is more important than competition (biotic stress) and that, under drought conditions, plants are more affected by water availability than by the competition with neighboring plants.

Plants have developed diverse strategies to mitigate stress, such as early flowering to “escape” abiotic stress and late flowering to mitigate biotic stress (Kigel *et al*., 2011). Plants growing naturally along environmental gradients provide a natural experiment for testing the hypothesis that natural selection led to local adaptation. Climatic gradients are especially useful in replacing space with time to detect adaptation to different climates (Etterson & Shaw, 2001). Nonetheless, climatic (abiotic) stresses are not exclusively affecting plants; biotic stresses, such as inter- and intraspecific competition also vary along the geographical scale and join abiotic stresses as selection agents (Seifan *et al*., 2010; Rysavy *et al*., 2014; Rysavy *et al*., 2016). Recent evidence claims that plants respond differently to single or to multiple simultaneous stresses (Rizhsky *et al*., 2004; Mittler, 2006; Mittler & Blumwald, 2010). While experimental studies usually test for the effect of a single stress, wild plant populations are usually exposed to a combination of biotic and abiotic stresses simultaneously (Ramegowda & Senthil-Kumar, 2015).

As already established in many previous studies (e.g., Franks *et al*., 2007; Franks & Hoffmann, 2011; Kigel *et al*., 2011; Penner *et al*., 2019), we also found that flowering time is an important phenotype for adaptation to climate, where plants originating from arid environments set flowering earlier in both experiments (Table 2; **Fig. 2a, 3b**). This result supports the hypothesis of escape strategy and indicates that plants from arid regions are preadapted to local abiotic stresses, whereas mitigation is achieved through early and rapid life cycle, allowing the plants to “escape” from the climatically difficult seasons (Ludlow, 1989; Kigel *et al*., 2011; Penner *et al*., 2019).

Current climate-change models for the Mediterranean climate region consistently predict an increase in aridity stress and, hence, a decrease in average water availability associated with increasing temperatures (Bates *et al*., 2008). In the eastern Mediterranean basin, annual precipitation is likely to significantly decrease while the temperature is expected to increase by the end of the 21^st^ century (Ben-Gai *et al*., 1998; Black, 2009; Black *et al*., 2010). Climate change is predicted to change species’ relative abundances and geographic ranges, causing extinctions if the plant response is not sufficiently rapid (Craine *et al*., 2011; Blois *et al*., 2013). Responses of plants to climate changes can be either short-term, such as phenotypic plasticity, or relatively slow, which relies on genetic variation and strong selection on heritable traits (Jump & Peñuelas, 2005; Barrett & Schluter, 2008; Anderson *et al*., 2012; Matuszewski *et al*., 2015). In this regard, the potential of a population to adapt to changes in climate will be at least partially governed by life history and phenology (Jump & Peñuelas, 2005). Notably, in our common garden experiments, we found that plants from along the entire aridity gradient grown in the arid research site flowered earlier than those grown in the Mediterranean site (Table 2, **Fig. 3b**), indicating a plastic response that enables plants from different climatic origins to change their life cycle rate according to the new conditions. Similarly, plants from the same origin were larger in the absence of stress (either abiotic or biotic) in comparison to plants under any of the stress treatments (**Fig. 2b, 3c**). This result also suggests a plastic response to stress.

Survival of plants also provided an indication of the role of multiple stresses in the evolution of adaptation. While in the controlled watering experiment all plants (100%) survived, in the arid common garden experiment survival was differential, where Mediterranean plants survived less than plants of an arid origin, and this phenomenon only occurred under competition (**Fig. 3a**). A high survival rate in the controlled watering experiment could result from a relative moderate Mediterranean climate that created a better environment, even with decreased watering. In the arid common garden, natural conditions are probably harsher and more unpredictable for plants of a Mediterranean origin, exposing them to stronger selection in early life stages. Although not tested or measured in this study, one can speculate about the role of high temperatures on survival probability in the arid site (Be’er Sheva), where aridity index is relatively low (Aridity index = 9.15) due to high mean temperature. Although the major change in the East Mediterranean is reduced precipitation, the combination of lower water availability with high temperatures and biotic (competition) stress may have a synergistic effect on plants (Black *et al*., 2010; Ramegowda & Senthil-Kumar, 2015; Rysavy *et al*., 2016).

Reproductive output, expressed here as the total number of spikes, is a fitness component that is directly associated with population dynamics and micro evolutionary change due to selection. If plants are locally adapted to their native habitat, they are expected to have higher fitness in their “home” environment, compared to “away” environments (Kawecki & Ebert, 2004). Here, we found opposing patterns of reproductive success in plants under dry and wet watering treatment (**Fig. 2c**): in the wet treatment, plants originating from arid regions had higher fitness than plants originating from Mediterranean regions, while this trend was reversed in the dry treatment – plants originating from arid regions had lower fitness than Mediterranean plants. These opposite trends may result from different resource allocation in the reproduction of plants originating from different environments, where plants from arid climate have a trade-off between rapid life cycle (“escape” strategy) and fitness. Local adaptation is driven by the accumulation of natural selection along many generations, providing a fitness advantage to plants in their native environments (Kawecki & Ebert, 2004; Dorman *et al*., 2009). We did not observe selection on flowering time in the controlled watering experiment (Table 4, **Fig. 4a**), while in the common garden experiments we found directional selection on flowering time (Table 4; **Fig. 4b, 4c**). As discussed above, the lack of selection observed in the controlled watering experiment may have resulted from the relatively benign conditions in this moderate Mediterranean climate, lacking other stress factors characterizing natural habitats. In contrast, the experimental common gardens were situated in natural conditions, and no artificial intervention was implemented after seed sowing. In the Mediterranean site (Tel Hai), competition treatment was the major stress affecting the direction of selection on flowering time (**Fig. 4b**), while in the arid site (Be’er Sheva), aridity stress was the major stress that eliminated the effect of competition on the selection on flowering time (**Fig. 4c**). Our phenotypic selection results from the common gardens suggest that the local adaptation of plants to their original habitats, through variation in flowering time, is a response to different stress factors. In the Mediterranean climate, where the main stress is biotic (competition), natural selection favors late flowering plants, but without competition, early flowering plants are favored. In the arid climate, where the main stresses are abiotic (water shortage), natural selection favors early flowering plants, regardless of competition, which is proposed to be lower than in the Mediterranean climate.

In summary, we found support for the hypothesis that flowering time is a key trait of annual plants in mitigating biotic and abiotic stresses. We also demonstrated that the local adaptation of plants to their climatic habitat is enabled due to variation in flowering time, as observed in a previous study (Penner *et al*., 2019), and that phenotypic selection is an important mechanism in the adaptation of plants to stress. While phenotypic adaptation through phenology is apparent, its genetic basis is still unknown. As a further study, we will quantify gene expression in *Brachypodium* plants from populations across aridity gradients in a controlled watering experiment to identify genes that are differentially expressed under aridity stress in populations that are adapted or not to aridity. The published *Brachypodium* genome (The International *Brachypodium* Initiative, 2010) is a valuable source that will enable annotation of these genes to provide the genetic basis for evolutionary mechanisms that are driven through natural selection.

## Acknowledgments

We thank S. Ezrati and A. Caicedo for providing plant accessions of the reference cytotypes. The watering experiment was logistically supported by TAUBG. We thank Prof. Kh. Kashkush and A. Dvir from Ben Gurion University of the Negev for their help in setting the Be’er Sheva research site, and to Dr. H. Shemesh from Tel Hai College for his help in setting this research site. M. Bar-Lev helped in phenotypic measurements at the Tel Hai site. R. Kent, G. Keren, E. Guk and O.-L. Har-Edom provided precipitation and temperature data and helped in producing the precipitation map. We thank K. Gallagher for English editing and insightful comments. We also thank Dr. U. Obolski for his useful statistical advises, and Dr. M. Seifan, Prof. I. Mayrose, Prof. Y. Ebenstein, Prof. L. Hadany and members of the Sapir lab for fruitful ideas and discussions throughout the course of this study.

## Supplementary captions

**Table S1** - Mean values (± S. E.) of phenotypic traits of *Brachypodium* from populations collected along the aridity gradient in Israel and included in the controlled watering experiment.

**Table S2** - Mean values (± S. E.) of phenotypic traits of *Brachypodium* from populations collected along the aridity gradient in Israel and included in the common garden experiment.

